# How do variable patch sizes hold higher beta diversity?

**DOI:** 10.64898/2026.09.29.755527

**Authors:** Minagi Naka, Akira S. Mori, Ryosuke Iritani

## Abstract

Beta diversity is a fundamental component of regional biodiversity that accumulates from compositional differences between local species assemblages. Although beta diversity is increasingly applied across spatial scales, its scale dependence remains incompletely synthesized, particularly concerning variation in spatial scale used to define alpha diversity (grain size). Using tree community data from Barro Colorado Island, we find that both additive and multiplicative beta diversity are consistently higher for unequal than equal grain divisions across different spatial extents and numbers of grains. These results can be explained by Jensen’s inequality based on the concave-down relationship of alpha diversity with area that in turn increases beta diversity through the scaling relationship between alpha and gamma diversity. Combined with our additional analysis showing that this prediction is robust to more general models of species richness, including species–area and species–sampling-effort curves, these results reveal a previously unrecognized, predictable range of beta diversity variation.

## 1 Introduction

Patterns of site-to-site variation in species composition, known as beta diversity, have long been a central topic in community ecology. Since beta diversity has the potential to improve predictions of natural and anthropogenic influences on biodiversity, many different measures of beta diversity have been introduced (Whittaker 1960; Baselga 2010; Anderson et al. 2011; Tatsumi et al. 2021). More recently, however, interest in beta diversity has returned to its original focus on scaling relationships between local (alpha) and regional (gamma) diversity (Whittaker 1972; Crist and Veech 2006; Chao et al. 2023). This shift reflects the growing recognition that beta diversity is a key component of regional biodiversity, associated with species distribution across space, regional buffering against local extinctions, and ecosystem stability and resilience. Consequently, beta diversity has become a central concept in conservation and restoration, where habitat heterogeneity is manipulated to maximize biodiversity at the landscape scale (Mori et al. 2018; Müller et al. 2023; Gonçalves-Souza et al. 2025). Yet, the assumptions underlying beta diversity measurements often differ between community ecology and applied contexts.

This issue is exemplified by the longstanding ‘single large or several small’ (SLOSS) debate within the conservation research community (Fahrig 2020; Fahrig et al. 2022). Traditionally, a single large patch of habitat holds more species than several small patches of the same total area (Diamond 1975). By contrast, recent evidence has demonstrated that small patches could enhance regional biodiversity through species replacement by explicitly integrating beta diversity components (Fahrig et al. 2022; Dambros et al. 2024; Perrin et al. 2025; Zhang et al. 2026). The key premise is the variation in the sizes of patches. Meanwhile, community ecologists recognize the spatial grain defining alpha diversity as a crucial attribute of beta diversity measurements (Barton et al. 2013; Spasojevic et al. 2016; Viana and Chase 2019). This scale dependence is likely because an increase in area leads to the accumulation of individuals and species and the averaging of spatial and environmental variation within plots, which subsequently influence beta diversity (Barton et al. 2013). Nevertheless, although beta diversity is known to scale with grain size, the impact of varying grain sizes within a single dataset remains poorly understood.

A possible explanation for the limited attention paid to the effects of unequal grain sizes on beta diversity is that the quantitative framework of beta diversity has been built upon the concept of alpha diversity (Whittaker 1960; Jost 2007; Anderson et al. 2011). Alpha diversity is inherently scale-dependent; species area relationships (SAR) have been studied as one of the most fundamental patterns in ecology, describing the tendency for larger areas to harbor a larger number of species across diverse taxa and habitats (Arrhenius 1921; Gleason 1922). Although the theoretical development of SAR is still a central topic in ecology (Borda-de-Água et al. 2025), numerous studies have shown that the number of species increases rapidly at small spatial scales and more gradually at intermediate area sizes, following a power-law relationship with an exponent less than one (Connor and McCoy 1979; Drakare et al. 2006; McGill 2011). Because the dependence of beta diversity on both alpha diversity and species pool size (i.e., gamma diversity) has long been recognized as the crux of beta diversity (Anderson et al. 2011; Chase et al. 2011; Chase and Myers 2011; Kraft et al. 2011), studies commonly use a standardized quadrat size to isolate the effects of spatial area from other variables. Such an evaluation approach, however, overlooks a fundamental mathematical artifact: whenever alpha diversity scales nonlinearly with area, the system’s average response to variable patch sizes systematically deviates from its response to average patch sizes.

This pervasive phenomenon is governed by Jensen’s inequality, named after the Danish mathematician Johan Jensen (Jensen 1906). The theorem formally demonstrates that variance in an explanatory variable inherently depresses the expected response of any concave function. In our context, because species-area relationships are universally concave in habitat area *x* (Preston 1962a; Preston 1962b; MacArthur and Wilson 1967; Coleman 1981), the mean local richness across variable habitat patches is strictly bounded above by the richness of a patch with the average area:

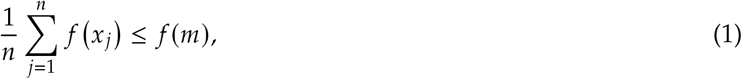

where *x*_*j*_ denotes the area of patch *j*, with *j* = 1, 2, … , *n*, and *m* = (*x*_1_ +…+*x*_*n*_)/*n* denotes the average area. Crucially, this inequality dictates that variance in patch sizes invariably depresses the average alpha diversity. Because beta diversity is fundamentally defined through the partition of a fixed regional species pool (gamma diversity) by this local richness, any mathematical deflation of alpha diversity must systematically inflate the resulting beta diversity. Consequently, unequal patch subdivision alone can generate an apparent increase in compositional heterogeneity, entirely as a mathematical artifact of non-linear averaging (Figure 1).

**Figure 1:**
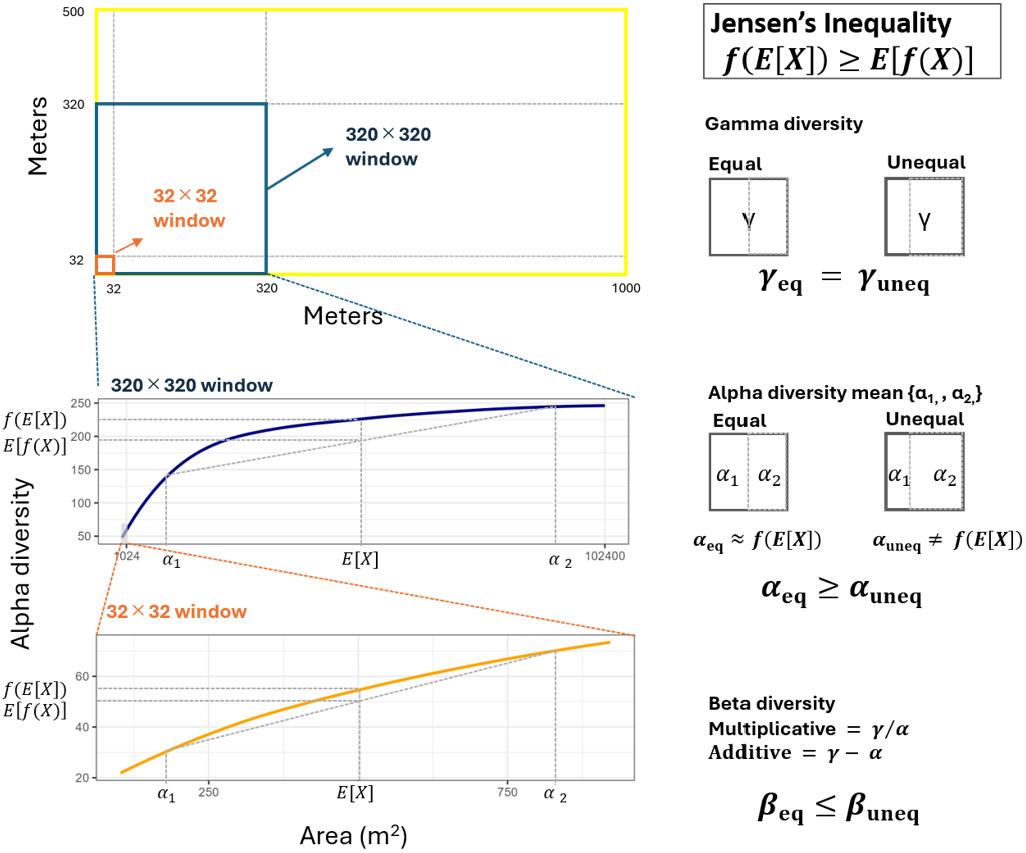
Schematic illustration of the sampling design. Tree alpha, beta, and gamma diversity were calculated across subsamples for each of two spatial extents shown in different colors. Blue lines indicate the 320× 320 m extent and orange lines indicate the 32 ×32 m extent. These extent windows were systematically moved across the 50-ha plot of Barro Colorado Island (BCI) in 20 m increments. Within each extent window, alpha diversity was calculated as the mean tree diversity across 2–256 subdivided sites, gamma diversity as the total tree diversity across all subdivided sites, and beta diversity as multiplicative (*β*_×_ = γ/ *α*) and additive (*β*_+_ = γ−*α*) beta diversity, representing variation in species composition among cells within each extent. According to our hypothesis, Jensen’s inequality predicts that variation in grain size reduces mean alpha diversity relative to the value expected from the average grain size. Because our sampling design manipulates only variation in local grain size while holding gamma diversity constant, both multiplicative and additive beta diversity are expected to increase as alpha diversity declines. We further hypothesized that the magnitude of the increase in beta diversity caused by variation in grain size would differ between spatial extents: it should be smaller within the 32m × 32m extent, where the species–area relationship (SAR) is nearly linear, and larger within the 320m × 320m extent, where the SAR is strongly concave.

Here, we connect the two cornerstones in ecology, species–area relationships and Jensen’s inequality, to investigate the effects of patch-size variation on beta diversity. We use two fundamental metrics, multiplicative and additive beta diversity, both of which are based on diversity-partitioning approaches, and formulate three hypotheses. First, the difference in beta diversity between systems with variable patch sizes and systems composed of equal-sized patches might increase with the concavity of the SAR. According to Jensen’s inequality, this deviation is expected to be larger at broader spatial scales, where the SAR is strongly concave, than at smaller spatial scales, where the SAR is nearly linear (Figure 1). Second, variation in patch sizes may determine this gap. Even when the total spatial area remains constant, the greater variation in patch size may result in higher beta diversity than that observed among equal-sized patches. Third, the effect should extend beyond pairwise comparisons and remain detectable when beta diversity is calculated across multiple patches. Throughout this paper, the term ‘grain size’ will refer to spatial scales defining alpha-diversity, which describes patch and local habitat.

## 2 Method and result

### 2.1 Site description

We illustrate our hypothesis using a data set of tree and shrub species from a rectangular 50ha plot (1000m × 500m) of old-growth tropical moist forest on Barro Colorado Island (BCI), Panama (Condit 1998; Hubbell et al. 1999; Condit et al. 2019a; Condit et al. 2019b). The BCI data set contains information on the spatial coordinates and diameter at breast height (dbh) of all woody plants (except for lianas) with stem diameter ≥ 1cm dbh, collected in different censuses. Since the results were similar across all censuses, we present results only for the 2005 census. The 2005 data set contains records on 301 species and 242,734 individual stems.

### 2.2 Grain set preparation and statistical analysis

We defined two spatial extents, namely a regional scale (320m × 320m) and a local scale (32m × 32m). To account for variation in beta diversity across different environmental conditions, regional and local windows were systematically slid across the 50-ha plot in 20-m increments. Because this procedure generated a larger number of windows at the local scale than at the regional scale, we randomly selected 350 local-scale windows to match the number of windows available at the regional scale. Then, each window was divided along both the horizontal and vertical axes into two subwindows with area ratios ranging from 1 : 9 to 9 : 1. To further evaluate the generality of Jensen’s inequality for regional beta diversity across varying spatial resolutions, we randomly partitioned each window into equal-sized grids of 2 × 2, 4 × 4, 8 × 8, and 16 × 16 grains. Multiplicative beta diversity (*β*_×_ = γ/*α*), and additive beta diversity (*β*_+_ = γ − *α*) (Whittaker 1960; Lande 1996; Veech et al. 2002b) were calculated for each set of grains generated by the different subdivision schemes within each window; we note that these two beta diversity metrics measure different aspects of spatial biodiversity: *β*_×_ represents the effective number of distinguishable local sites, whereas *β*_+_ represents the effective amount of diversity gained by expanding the sampling scale from a single average site to the entire region. Species area relationships were confirmed on regional and local scales. For each spatial extent and beta diversity metric, differences in beta diversity among the different subdivision ratios were assessed using multiple-comparison tests, with the glht function of the ‘multcomp’ package for multiple comparisons with Tukey’s test. For grain numbers greater than two (2 × 2, 4 × 4, 8 × 8, and 16 × 16), beta diversity obtained from randomly subdivided grains was compared with that obtained from equally spaced subdivisions using *t*-tests, with the t.test function of the ‘dplyr’ package.

### 2.3 Key result

Both multiplicative and additive beta diversity were higher under varying grain size ratios than under equal grain sizes at the local and regional scales. These deviations increased as differences in grain size ratio increased (Figure 2a-d). For additive beta diversity, the mean value for the 1 : 9 grain size ratio exceeded that for the equal-size ratio by 18.6 at the regional scale and by 7.8 at the local scale. For multiplicative beta diversity, the corresponding differences were 0.10 and 0.23, respectively. When the number of grains exceeded two, beta diversity was consistently higher in randomly than equally subdivided grain sets (Figure 3).

**Figure 2:**
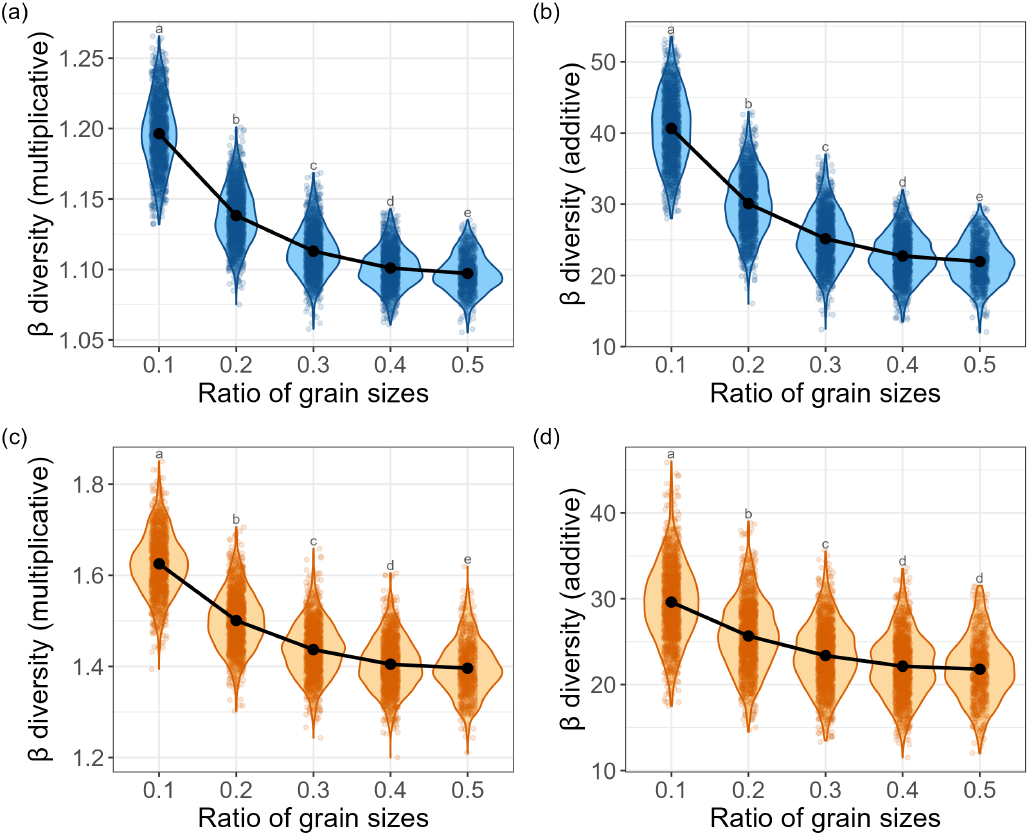
Changes in beta diversity across different ratios of two spatial grains: (a) multiplicative beta diversity at the regional scale, (b) additive beta diversity at the regional scale, (c) multiplicative beta diversity at the local scale, and (d) additive beta diversity at the local scale.

**Figure 3:**
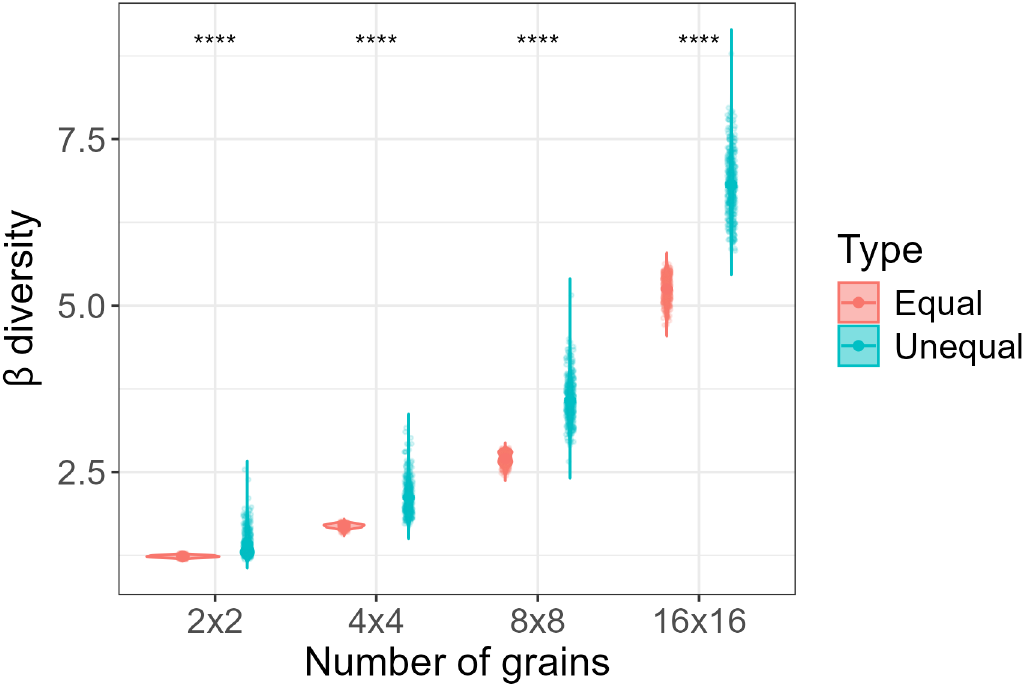
Changes in beta diversity across different numbers of grains. Red and blue dots represent multiplicative beta diversity calculated among equal-sized grains and among randomly subdivided grains of different sizes, respectively.

To analytically estimate the deviations, we write the deviations of alpha and beta diversity as:

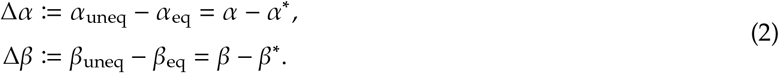

We refer to these deviations as ‘Jensen gaps’ in the present manuscript. To approximate the Jensen gaps, we use the delta method (Cox 2005) for alpha diversity:

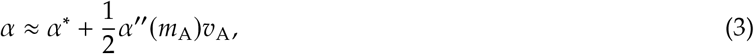

where the double prime represents the second derivative with respect to the area, evaluated at the equal partitioning (which will apply henceforth unless otherwise stated). Here, *m*_A_ and *v*_A_ represent the average and variance among grain sizes *A*, respectively. The second term in Equation (3) is negative whenever species richness *α*(*A*) is concave in grain size *A* (e.g., *α* = *cA*^*z*^, with 0 < *z* < 1, as in the standard species–area curve). As a result, the Jensen gap in alpha diversity, Δ*α*, is approximated as:

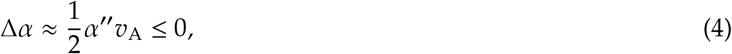

which is always nonpositive, with the equality for the equipartition case. Accordingly, the deviation in additive beta diversity Δ*β*_+_ is approximated as:

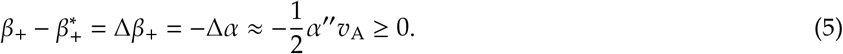

Equation (5) tells us that the variance in grain sizes *v*_A_ increases beta diversity. In other words, the equipartition minimizes beta diversity.

Similarly, the deviation in multiplicative beta diversity Δ*β*_×_ is given by:

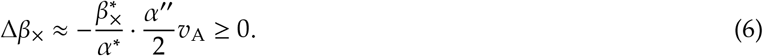

Again, the deviation in beta is proportional to the variance in grain sizes, but is now multiplied by an additional term due to the nonlinearity of beta. Consequently, the Jensen gap is positive whenever the grain size has a variation.

These approximations highlight the critical need to account for grain-size variance when interpreting beta diversity (especially in conservation strategies such as the SLOSS debate, where unequal patch sizes are prevalent). We provide more accurate approximations in Appendix A.

Based on the delta method, we predicted beta diversity in systems with randomly sized grains in comparison to beta diversity from an equivalent system of equally sized grains. Although the predicted values generally fell within the range of field observations, the prediction accuracy of the Jensen gaps depended on the assumed pecies area relationships (SAR) exponent (*z*; Figure 4). Predictions based on *z* = 0.25 were most accurate when the window was divided into 16 × 16 grains, whereas those based on *z* = 0.50 performed better for the remaining grain-size configurations.

**Figure 4:**
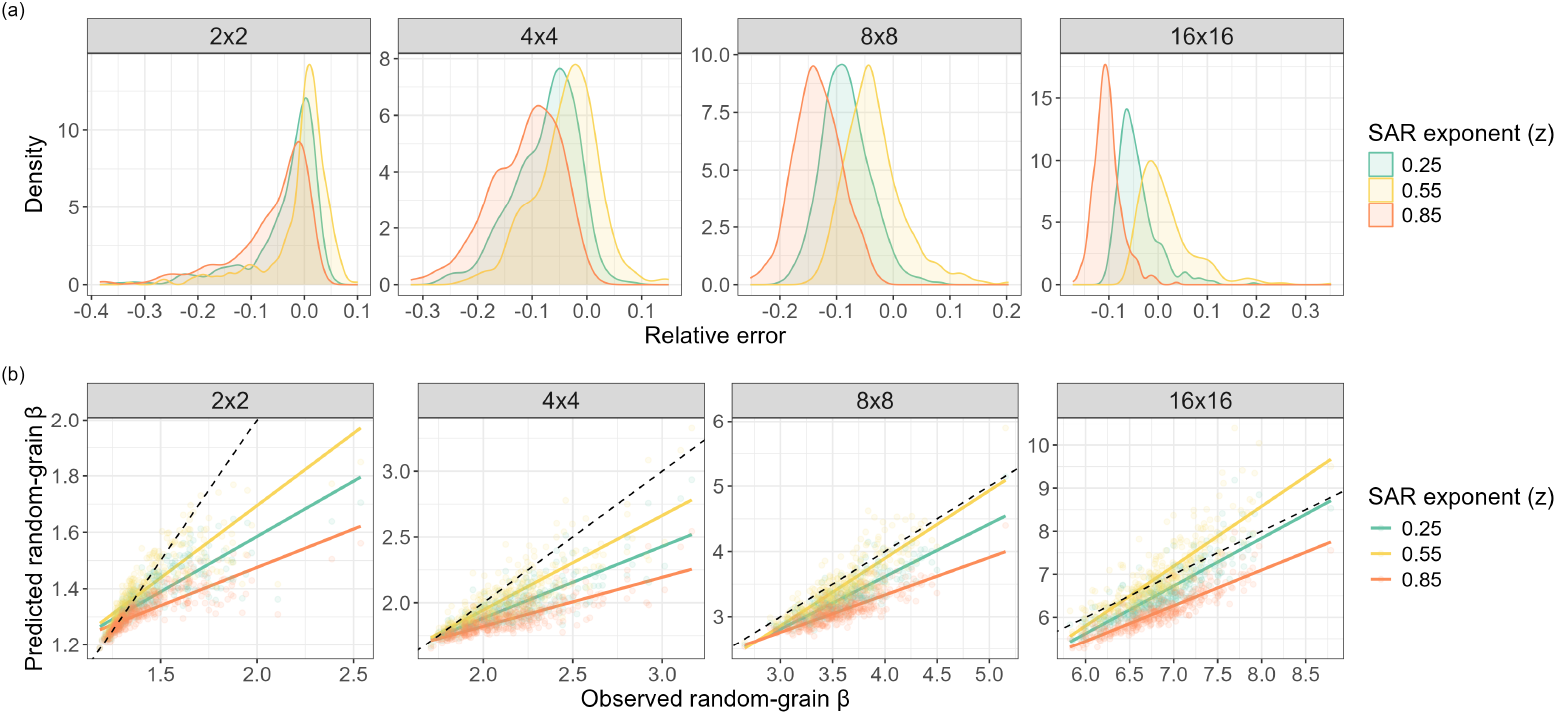
Sensitivity analysis of the Jensen gap approximation for predicting multiplicative beta diversity across randomly sized grains. (a) Distribution of relative error between predicted and observed beta diversity and (b) relationships between predicted and observed beta diversity across different numbers of grains. Predicted beta diversity was calculated using the delta method with three species–area relationships (SAR) exponents (z = 0.25, 0.55, 0.85). Colors represent different values of the SAR exponent (z).

Since the dependence of multiplicative beta diversity on alpha diversity is complex, our main results focus on the multiplicative beta diversity at the regional scale for quantifying beta diversity across multiple patches, and its sensitivity analysis (Figures 3 and 4). We additionally analyzed additive beta diversity in the Supplementary Information and obtained similar results (Figures S2 and S3). Overall, our additional analyses demonstrated the robustness of our findings to the choice of beta diversity metric - either additive or multiplicative.

### 2.4 Extension

So far, our analytical framework has relied on deterministic species–area relationships and natural settings where environmental heterogeneity has been confirmed to be less influential. To evaluate the robustness of these findings amidst natural ecological complexity, we generalized our predictions in two directions (Appendix B). First, we replaced the specific functional form of the SAR with the broader mathematical concept of submodularity. Ecologically, submodularity formalizes the principle of diminishing returns in species discovery. Intuitively, as a sampled habitat expands, the rate of encountering novel species naturally declines due to an increasing spatial overlap of species compositions. By combining this intrinsic biological property with majorization theory and an analytical refinement for the Jensen gap (Liao and Berg 2019), we proved that unequal habitat partitioning strictly reduces expected alpha diversity, entirely independent of the underlying spatial configuration. Second, we extended our framework to a fully probabilistic stochastic landscape model (Iritani et al. 2025). This confirmed that the inflation of beta diversity under uneven sampling is a robust, universal consequence of these diminishing returns in species accumulation, beyond the standard species-area curves such as *cA*^*z*^ (Darlington 1943; Preston 1962a; Preston 1962b; MacArthur and Wilson 1967). Consequently, our predictions remain highly robust across a broad class of realistic species–area relationships.

## 3 Discussion

This study uncovered the underlying principle governing the effects of grain size variation on beta diversity. Under conventional equal-weight diversity partitioning, consistent with our first hypothesis, a concave species–area relationship (SAR) leads to a predictable increase in beta diversity with unequal grain sizes compared with equal grain sizes (Figure 2). Notably, the magnitude of this effect can then be approximated from the curvature of the SAR and the variance in grain size, as predicted by the second hypothesis (Figure 4). Furthermore, the consistency of these results across multiple patches supports our third hypothesis (Figures 3 and 4). This knowledge would allow for disentangling variation in beta diversity driven by biological dynamics that are induced by differences in grain size. The importance of our findings may be further underscored by growing conservation and restoration efforts to maximize landscape biodiversity through the manipulation of habitat sizes within a region and, ultimately, beta diversity. Incorporating the perspective of beta diversity into the applied fields sheds new light on questions of beta diversity.

Using the tree community dataset from Barro Colorado Island (BCI) , we confirmed that beta diversity was higher when grain sizes varied than when grain sizes were equal. Although multiple drivers jointly influence patterns of beta diversity, these differences can be accounted for by Jensen’s inequality as a consequence of the averaging of alpha diversity. Specifically, if alpha diversity changes nonlinearly with area, as observed in this study (Figure 1) and consistent with established theory (Hubbell 2001; Borda-de-Água et al. 2025), Jensen’s inequality explains that increasing the variance at grain size should reduce the mean value of alpha diversity. We find that increasing the variation in local grain size while keeping gamma diversity fixed leads to consistent increases in both multiplicative and additive beta diversity. Nonlinear SAR has been widely documented across a range of climates and taxonomic groups. For example, Watson 1859 first reported a convex relationship between species richness and area in plant communities on the island of Great Britain, and MacArthur and Wilson 1967 subsequently observed convex species–area relationships for reptiles and amphibians in the West Indies. Therefore, our results suggest that such scaling effects are likely inherent to many ecological systems.

By comparing local and regional scales, our findings further demonstrated that a substantial portion of the observed variability in beta diversity can be explained by Jensen’s inequality rather than by differences in underlying biological processes such as dispersal limitation and species interactions. For instance, the Jensen gap of additive beta diversity was smaller at local scales, where the SAR was nearly linear, than at regional scales, at which the SAR was strongly concave (Figure 2bd). These results suggest that the shape of the SAR may determine the magnitude of effect of unequal partition. In contrast, this effect was smaller for multiplicative beta diversity at the regional scale (Figure 2ac). These results suggest that the Jensen gap of multiplicative beta diversity does not depend only on alpha but also on gamma diversity. Additionally, while the present study focused on the spatial scale where the SAR shows a slower growth rate with area, SAR exhibits rapid growth again at the continental scale in several systems (Hubbell 2001; McGill 2011). Based on Jensen’s inequality, average alpha diversity is expected to be higher in systems with variable patch sizes than in systems composed of equal-sized patches when the SAR is convex. Although further investigation for the shape of SAR is needed, our results hypothesize that higher beta diversity may be observed among the equal-sized patches at transition zones between the regional and continental scales.

We found that the variation in grain size strongly influences the Jensen gap, consistent with previous studies applying nonlinear averaging to biological patterns (Ruel and Ayres 1999; Denny 2017). While these results may seem to highlight the context-dependency of the Jensen gap, they can be predicted to a considerable extent using the delta method (Equations (3), (5) and (6)). Importantly, even when there are multiple grains, predictable differences in beta diversity may still exist (Figure 4). Accordingly, interpreting beta diversity requires considering grain size variation based on our finding that higher beta diversity was consistently observed for randomly sized grains regardless of the number of grains (Figure 3). We further generalized this beta-diversity pattern by demonstrating that it applies to any gradient in which alpha diversity changes nonlinearly, as described by a submodular function (e.g., species–area relationship, or species–sampling effort relationship; Appendix B). This is particularly relevant in applied conservation contexts, where scaling relationships between local and regional diversity are often compared among patches of unequal size, or unequal sampling effort (Fahrig 2020; Dambros et al. 2024). For example, Fahrig (2020) reviewed 157 habitat fragmentation studies and found that more than half were potentially affected by sampling bias, such as a decline in sampling intensity with increasing patch size (despite attempts to compensate for these biases). Our theoretical framework may therefore facilitate more robust predictions of beta diversity when habitat area and sampling bias are inherently confounded.

Our findings also might be regarded as a consequence of beta diversity arising from the potential pitfall of averaging alpha diversity across heterogeneous spatial grains. However, we argue that these results should be understood from the beta diversity perspective, in particular for two reasons. First, averaging alpha diversity is used for two classical formulations of beta diversity, multiplicative and additive partitions (Whittaker 1972; Lande 1996; Veech et al. 2002a). Indeed, explicit dependence on mean alpha diversity is restricted to diversity-partitioning approaches, including these two beta diversity indices (Baselga 2010; Anderson et al. 2011). Still, these indices have important implications for management strategies aimed at maximizing biodiversity at the landscape scale (Socolar et al. 2016). For instance, additive partitioning quantifies the number of species added by beta diversity to the regional diversity, whereas multiplicative partitioning quantifies the relative contribution of beta diversity to regional diversity (Dambros et al. 2024). In light of the renewed recognition of their utility in applied contexts and the development of new beta diversity measures, such as Hill-number-based diversity partitioning, these patterns should be interpreted through the lens of beta diversity (Hill 1973; Socolar et al. 2016; Mori et al. 2018; Chao et al. 2023; Müller et al. 2023).

Second, long-standing debates such as SLOSS, which concern how habitat area should be allocated while keeping total area constant, ultimately center on the extent to which beta diversity can be manipulated (Fahrig 2003; Fahrig 2020). Because these issues consider patch size and patch number jointly, beta diversity has typically been evaluated with little attention to variation in patch size (Dambros et al. 2024; Gonçalves-Souza et al. 2025; Zhang et al. 2026). Our findings, however, suggest that the contribution of beta diversity should be evaluated more cautiously, as beta diversity is likely to be overestimated when grain size and its variance are not considered, as may occur in meta-analyses. Conversely, if observed patterns are inconsistent with our prediction (e.g., higher beta diversity is observed among equal-sized grains in systems with a concave SAR), this would provide stronger evidence that the observed beta diversity reflects ecological processes rather than the scaling effect described here. Importantly, inequality is not an ecological phenomenon per se, but instead a mathematical consequence of the nonlinear form of many biological functions and the variance inherent in many environmental factors. Despite its potential importance, the implications of the Jensen gap remain incompletely understood and warrant further investigation.

We remark that the possible bias in our findings may arise from spatial autocorrelation. Indeed, our sensitivity analysis shows that predicted beta diversity with unequal grain sizes from that with equal grain sizes, tended to be lower than the observed values (Figure 4). Although spatial aggregation patterns of species can vary across geographical locations and spatial scales (i.e., spatial extents and grain sizes; Spasojevic et al. 2016; Nishizawa et al. 2022), failing to account for spatial autocorrelation likely underestimates the difference in beta diversity between unequal and equal grain sizes (but see Appendix B for a formal framework to explicitly isolate the effect of spatial covariance). Additionally, our analysis was based on a continuous tree census plot, whereas habitat fragmentation, as discussed in the SLOSS debate, is often influenced by various factors such as edge effects, the spatial arrangement of habitats, and surrounding land-use types. Therefore, further studies will be needed when extrapolating these findings to landscape-level biodiversity conservation.

## 4 Conclusion

This study tackled the potential generality of changes in beta diversity caused by grain size variations by integrating pecies area relationships (SAR) with Jensen’s inequality, two well-established frameworks in ecology. We found that beta diversity was consistently higher among unequally divided grains than among equally divided grains for additive and multiplicative beta diversity metrics, regardless of spatial extent or number of grains. Moreover, our theoretical extension showed that the observed difference includes a purely mathematical effect whose magnitude can be estimated from variation in grain size. Our results therefore indicate that accounting for the effects of habitat size variation on beta diversity could help disentangle variation in beta diversity driven by other factors, including environmental heterogeneity, disturbance, and biological interactions. Assessing the spatial dependence of beta diversity will provide deeper insights into how biodiversity is spatially organized, the central question in any field of ecology.

## 5 Acknowledgements

We sincerely thank David Alonso and William Godsoe for helpful comments on this manuscript. The BCI forest dynamics research project was made possible by National Science Foundation grants to Stephen P. Hubbell: DEB-0640386, DEB-0425651, DEB-0346488, DEB-0129874, DEB-00753102, DEB-9909347, DEB-9615226, DEB-9615226, DEB-9405933, DEB-9221033, DEB-9100058, DEB-8906869, DEB-8605042, DEB-8206992, DEB-7922197, support from the Forest Global Earth Observatory (ForestGEO), the Smithsonian Tropical Research Institute, the John D. and Catherine T. MacArthur Foundation, the Mellon Foundation, the Small World Institute Fund, and numerous private individuals, and through the hard work of over 100 people from 10 countries over the past three decades. The plot project is part the Forest Global Earth Observatory (ForestGEO), a global network of large-scale demographic tree plots. This work was supported by JSPS Research Fellowships for Young Scientists.

## A Derivations of the approximations

### A.1 Alpha

We partition the region (with fixed area *A*_T_) into *n* subpopulations as {*A*_1_, … , *A*_*n*_} where 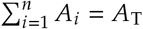 and *A*_*i*_ = *A*_T_ and *A*_*i*_ > 0, for *i* = 1, 2, … , *n*.^1^ We write *f* (*A*) for the species richness of a habitat of area *A*. Alpha diversity is the arithmetic mean of the local richnesses:

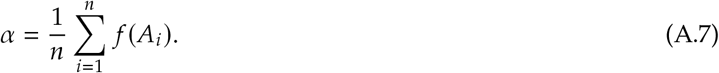

When all grain sizes are equal, we have:

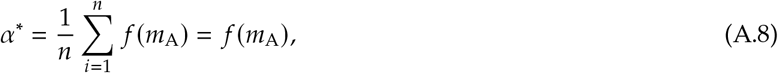

where *m*_A_ is the average grain size. Assuming that *f* is smooth (i.e., differentiable), we expand *f* (*A*_*i*_) around the average grain size *m*_A_:

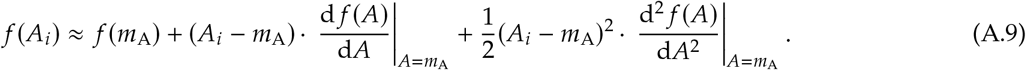

Taking the arithmetic mean over *i* = 1, 2, … , *n* of both sides, we get:

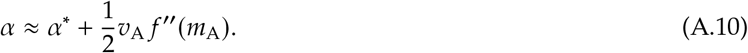

Because *f* ^′′^(*m*_A_) = *α*^′′^(*m*_A_), we recover the formula of the deviation in alpha diversity of the main text.

### A.2 Additive beta

The result of the additive beta diversity is straightforward since γ − *α* = *β*_+_ with γ fixed. Hence,

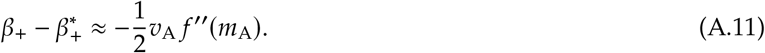

### A.3 Multiplicative beta

The multiplicative beta diversity *β*_×_ is defined by:

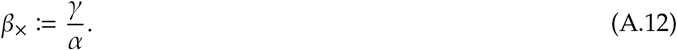

Because γ is fixed by the total area *A*_T_ and is unaffected by how the region is partitioned, the deviation 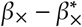 can be obtained by direct algebra, without any approximation:

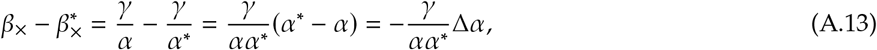

where Δ*α* = *α*−*α*^∗^ as in the main text. To express Equation (A.13) in terms of the grain-size variance *v*_A_, we substitute the delta-method approximation for Δ*α* (Equation (A.10)). Because Δ*α* = *O*(*v*_A_), we have *α* ≈ *α*^∗^ to leading order, so 1/(*αα*^∗^) ≈ 1/(*α*^∗^)^2^ up to an error of order *v*_A_; substituting yields:

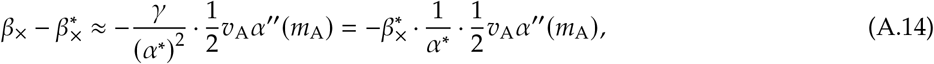

which recovers Eq. (6) of the main tex

### A.4 More accurate result

Let *A* be a discrete random variable representing the area of a subpopulation, uniformly drawn from {*A*_1_, … , *A*_*n*_}. This is precisely the finite-sample setting of Corollary 1 in Liao and Berg (2019), which specializes their Theorem 1 to an empirical distribution on *n* data points. For an open interval (*a, b*) ⊂ℝ, let ϕ : (*a, b*) → ℝ be a twice differentiable function. Liao and Berg (2019, Theorem 1) show that for any random variable *A* with *P*(*A* ∈ (*a, b*)) = 1 and mean *m*_A_,

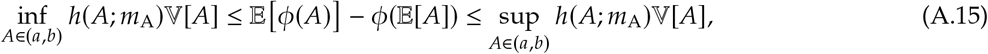

where:

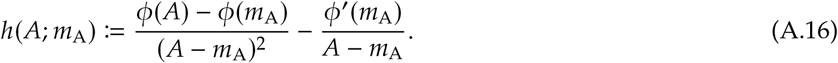

Because *A* here is uniformly drawn from the observed grain sizes {*A*_1_, … , *A*_*n*_}, V[*A*] = *v*_A_ is exactly the empirical grain-size variance defined above, and this bound is equivalently obtained from Liao and Berg (2019, Corollary 1).^2^

In our context, since the species–area relationship *f* (*A*) is concave, we apply the theorem to the convex function ϕ(*A*) ≔ − *f* (*A*). The Jensen gap is then E [−*f* (*A*)] −(−*f* (*m*_A_)) = *α*^∗^ − *α*. Computing *h*(*A*; *m*_A_) for − *f* (*A*) yields:

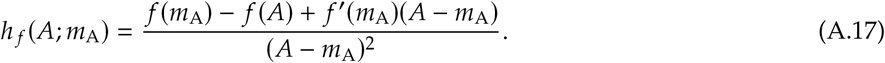

Because the deviation in alpha diversity is Δ*α* = *α* − *α*^∗^, we obtain bounds for the deviation of additive beta diversity (Δ*β*_+_ = −Δ*α*):

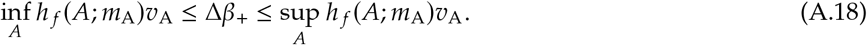

Furthermore, recall from Equation (A.13) that the exact deviation for multiplicative beta diversity is 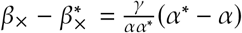. Since *α*_min_ ≤ *α* ≤ *α*_max_, substituting the bounds for *α*^∗^ − *α* yields the absolute bounds for the deviation:

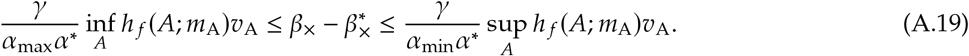

This mathematical result proves that as long as there is spatial variance in grain sizes (*v*_A_ > 0), the deviation of beta diversity is strictly positive and bounded by the intrinsic heterogeneity of the landscape. When the richness follows the power-law *α*(*A*) = *cA*^*z*^ (where 0 < *z* < 1), we can analytically specify these bounds. The auxiliary function becomes:

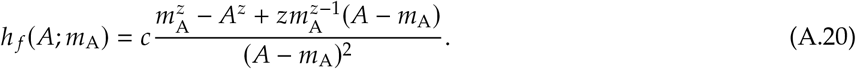

Because the third derivative of − *f* (*A*) is strictly negative (*cz*(1 − *z*)(*z* − 2)*A*^*z*−3^ < 0), its first derivative, − *f* ^′^(*A*), is a concave function of *A*. By Liao and Berg (2019, Lemma 1) — if ϕ^′^(*x*) is concave then *h*(*x*; *m*) is monotonically decreasing in *x* — this concavity guarantees that *h* _*f*_ (*A*; *m*_A_) is monotonically decreasing with respect to *A*. Therefore, if the unequal grain sizes fall within a specific range [*A*_min_, *A*_max_], the infimum and supremum are exactly attained at the boundaries. Substituting these limits yields the exact bounds for the deviation of additive beta diversity:

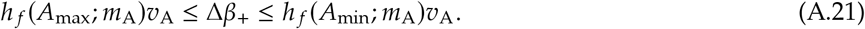

Similarly, for the multiplicative beta diversity, the rigorous bounds are explicitly given by:

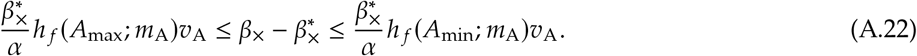

This formulation demonstrates how the degree of nonlinearity (governed by the exponent *z*) and the spatial bounds of grain sizes (*A*_min_ and *A*_max_) jointly and strictly constrain the scale of beta diversity deviation.

## B Extensions

### B.1 Setup

We consider the smallest grids and name them as Ω = {*g*_1_, … , *g*_*N*_} , where Ω is the region and *N* the number of grids; typically *N* = 4^*m*^, for some integer *m*. We call *g*_*i*_ as the *i*-th microsite (or simply microsite) for *i* = 1, 2, … , *N*. We say *G* is a subcommunity if *G* is nonempty and can be written as the union of microsites; for example, *G* = {*g*_1_} ∪{*g*_2_} = {*g*_1_, *g*_2_}. In other words, *G* is a subcommunity if *G* is a nonempty subset of Ω. We write #*G* for the number of microsites belonging to a subcommunity *G*. For example, # {*g*_1_, *g*_2_, *g*_10_} = 3. We say *k* = #*G* is a size of a subcommunity *G*. For a collection of nonempty subcommunities **G** ≔ {*G*_1_, … , *G*_*n*_}, we say **G** is an *n*-partition of the region if:

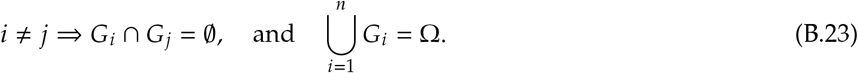

For an *n*-partition of the region, **G**, we write **k** = # ○ **G** ≔ (*k*_1_, … , *k*_*n*_) ≔ (#*G*_1_, … , #*G*_*n*_) (the element-wise cardinality) for the sizes of the subcommunities associated with **G**. Note that 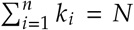 and and *k*_*i*_ ≥ 1, *i* = 1, 2, … , *n*. When *N* = *ne* for some integer *e*, we write **e**^∗^ ≔ (*e* , … , *e*) for the partition of the region size into the same size, or we call this as the *n*-equipartition.

### B.2 Richness accumulation function (SAF)

We write *r* : 2^Ω^ → Z for the species richness *r*(*G*) in a subpopulation *G*; we refer to *r* as the richness accumulation function. For example, when Ω = {*g*_1_, *g*_2_, *g*_3_, *g*_4_} and if we take *G* = {*g*_1_, *g*_3_} ∈ 2^Ω^, *r*(*G*) represents the richness of the subpopulation that pools two microsites *g*_1_ and *g*_3_.

### B.3 SAF is submodular

We observe that SAF is submodular: we pick two disjoint subcommunities *G* and *G*^′^, and consider adding one microsite *g* ∉ *H* ≔ *G* ∪ *G*^′^; then

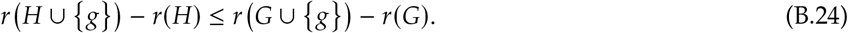

Biologically, this inequality reflects diminishing returns in species discovery. The number of newly encountered species in *g* is generally smaller for *H* = *G*∪*G*^′^ than for *G* alone, due to a higher likelihood of species overlap between {*g*} and *G*^′^ (Figure S1). When the species richness can be predicted by the area of a subpopulation, Equation (B.24) represents the concavity of species–area relationship. Therefore, the submodularity approach is a generalization of the classical species–area curve.

**Figure S1:**
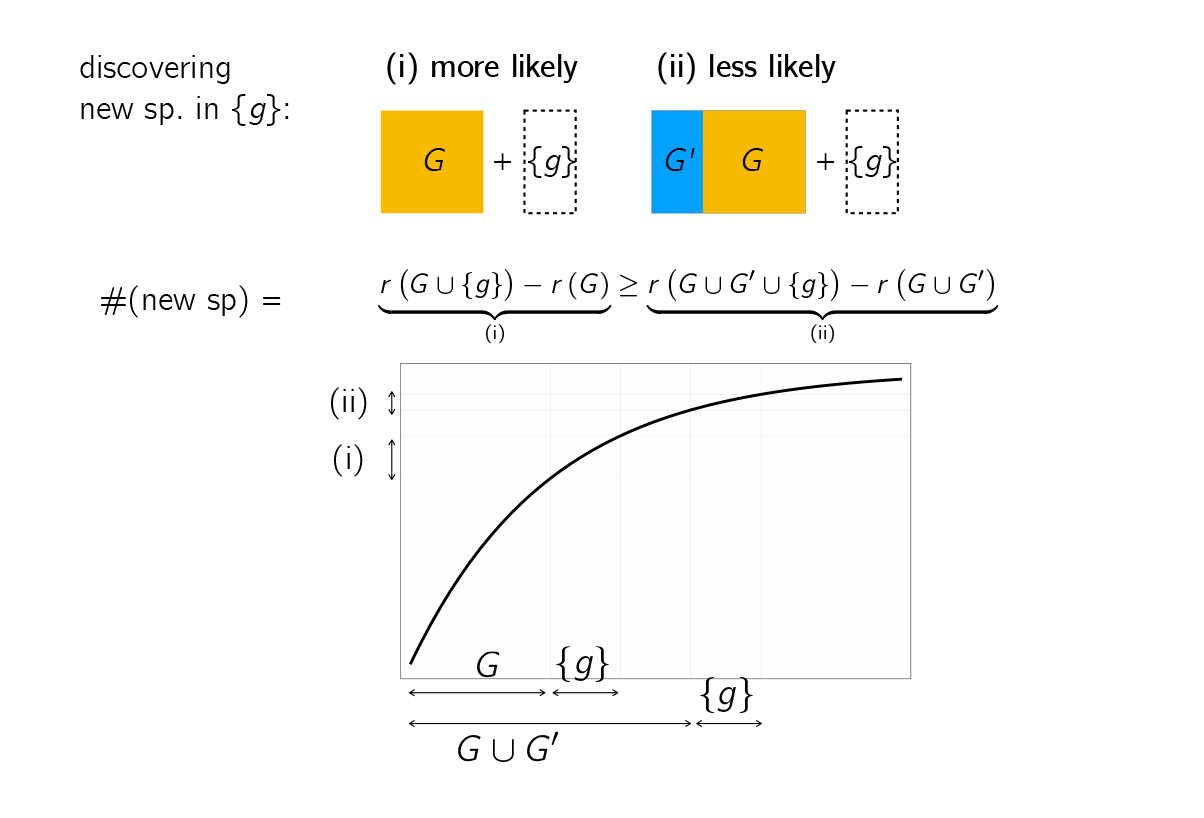
Conceptual representation of submodularity. The marginal gain in species richness from adding a local habitat patch (*g*) diminishes as the background community expands from *G* to *G* ∪ *G*^′^. In the context of the SAR, adding {*g*} to the census corresponds to including the previously uncensused area of *g*. This property of diminishing returns generalizes the concavity of the SAR to arbitrary spatial configurations.

### B.4 Probabilistic approach

Suppose we uniformly at random sample a subset of *k* micorsites to form a subpopulation *G* with #*G* = *k*. We define the expected species richness of a subcommunity *G* of size *k*, denoted by *R*(*k*), by taking the expectation over all possible subsets sized *k*:

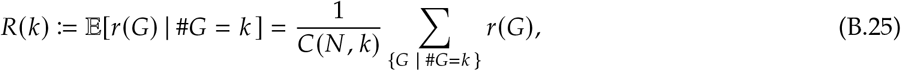

where *C* represents the binomial coefficient. We write Δ*R*(*k*) ≔ *R*(*k* + 1) − *R*(*k*) for the marginal gain in species richness obtained by sampling one additional microsite. We uniformly at random sample a subcommunity *G* of size *k* − 1 and two microsites {*g*} and {*g*^′^} with {*g, g*^′^} ∩ *G* = ∅. Using Equation (B.24) yields:

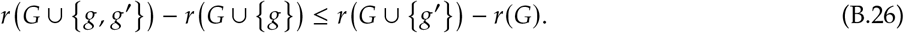

Taking the expectation in both sides in Equation (B.26), we get:

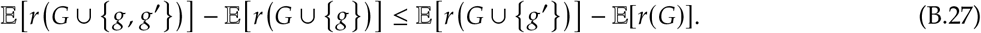

Since #*G* = *k* − 1, the left hand side is *R*(*k* + 1) − *R*(*k*) = Δ*R*(*k*) and the right hand side is *R*(*k*) − *R*(*k* − 1) = Δ*R*(*k* − 1). Therefore, Δ*R*(*k*) ≤ Δ*R*(*k* − 1), which means Δ*R*(*k*) is decreasing in the grain size *k*. In other words, the expected species accumulation function *R*(*k*) is discrete concave in the subcommunity size *k*.

### B.5 The equipartition maximizes alpha diversity

For a partition **G** = {*G*_1_, … , *G*_*n*_} with **k** = # ○ **G**, alpha diversity is the arithmetic average of local species richnesses:

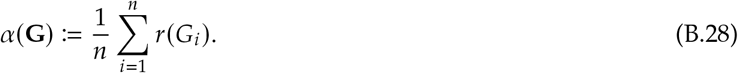

Fixing an *n*-partition size **k**, we can take the expectation of all possible partitions {**G**} such that # ○ **G** = **k**:

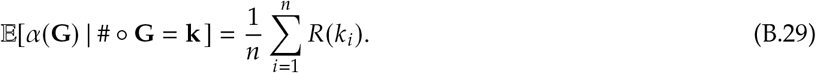

To show that the *n*-equipartition maximizes the alpha diversity, we use the theory of majorization (Marshall et al. 1979). This is a direct consequence of Karamata’s inequality, implying:

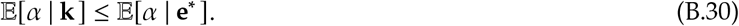

### B.6 The equipartition minimizes additive beta diversity

Using the same argument as in the main text, we can show that the additive beta diversity is minimized for the equipartition.

### B.7 The effect of the equipartition on multiplicative beta diversity

It is more difficult to assess the effect of the equipartition on the multiplicative beta diversity, because the alpha diversity appears in the denominator: *β*_×_ = γ/*α*. Unlike the additive case, evaluating E[*β*_×_ | **k**] exactly requires the expectation of a ratio, which is analytically intractable in general. We therefore combine the discrete-concavity result above with a second-order (delta-method) approximation, and are explicit below about which part of the resulting expression is fully justified by submodularity alone and which part is not.

Let δ_*i*_ = *k*_*i*_ −*e*^∗^ be the deviation of the *i*-th subcommunity size from the equipartition size *e*^∗^, such that 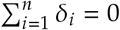. We define the variance of patch sizes (the degree of unequal partitioning) as 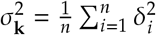. Expanding the expected species accumulation function *R*(*k*_*i*_) around *e*^∗^, we obtain:

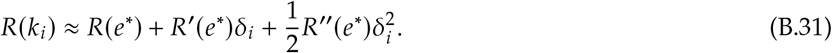

Taking the arithmetic mean over all *n* subcommunities, the first-order term cancels out because 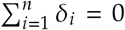. The expected alpha diversity under the partition **k** is then given by:

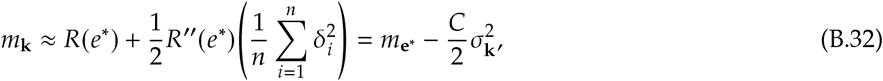

where *C* = −*R*^′′^(*e*^∗^) > 0 due to the discrete concavity (submodularity) of *R*. This equation explicitly demonstrates that the reduction in mean alpha diversity is directly proportional to the variance of the patch sizes.

Applying the delta method to the ratio γ/*α* (i.e., a second-order Taylor expansion of *g*(*x*) = γ/*x* around *m*_**k**_ = E[*α* | **k**], using *g*^′′^(*x*) = 2γ/*x*^3^) gives the standard approximation

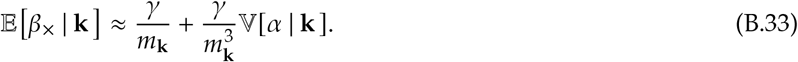

Comparing this to the equipartition value and substituting Equation (B.32) together with the approximation (1 − *x*)^−1^ ≈ 1 + *x*, the first term expands, to leading order in 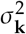, as

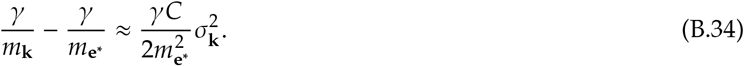

This term is fully determined by the discrete concavity of *R* (i.e., by *C* = −*R*^′′^(*e*^∗^) > 0) and mirrors, at the level of expected values, the mean-reduction mechanism already established for additive beta diversity: uneven partitioning lowers mean alpha diversity, which mechanically inflates γ/ *α*.

A second contribution comes from the variance term, 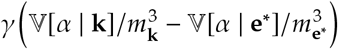 . Unlike Equation (B.34), this term is not pinned down by submodularity alone: it depends on how the sampling variance of *α*, V[*α* | **k**], changes as the partition becomes more heterogeneous, which in turn depends on the covariance structure among the richnesses *r*(*G*_*i*_) of differently sized subcommunities (and, in an applied setting, on spatial autocorrelation; cf. the discussion in the main text). Characterizing this dependence in general requires further assumptions that we do not attempt to justify here. We therefore report the two contributions separately rather than collapsing them into a single closed-form expression in 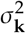:

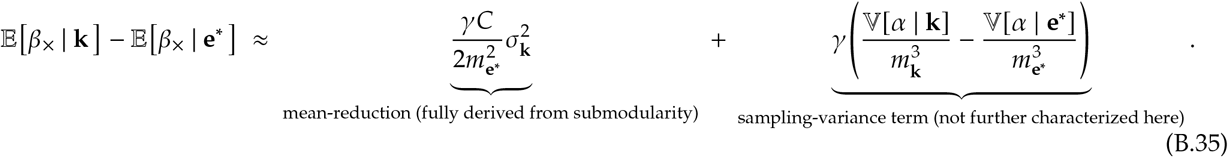

The first, mean-reduction term shows that even absent any change in sampling variance, the discrete concavity of *R* alone is sufficient to inflate the expected multiplicative beta diversity under unequal partitioning, in proportion to 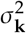. Whether the second term adds to or partly offsets this effect in a given system is an empirical question that depends on the spatial structure of the data, and we leave a general theoretical treatment of this term to future work.

**Figure S2:**
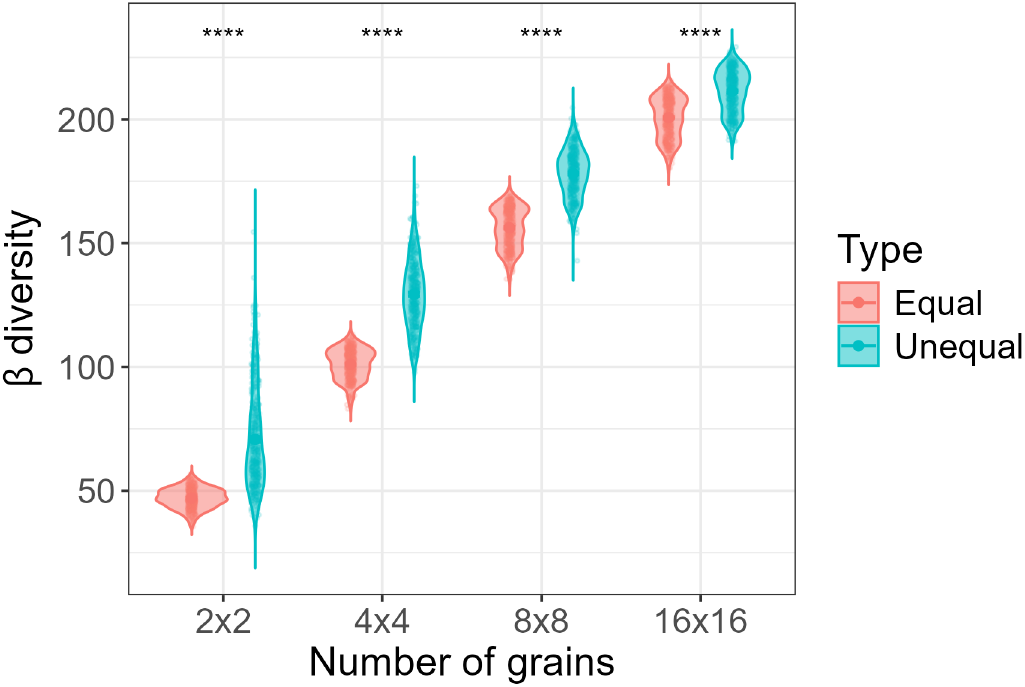
Changes in beta diversity across different numbers of grains. Red and blue dots represent additive beta diversity calculated among equal-sized grains and among randomly subdivided grains of different sizes, respectively.

**Figure S3:**
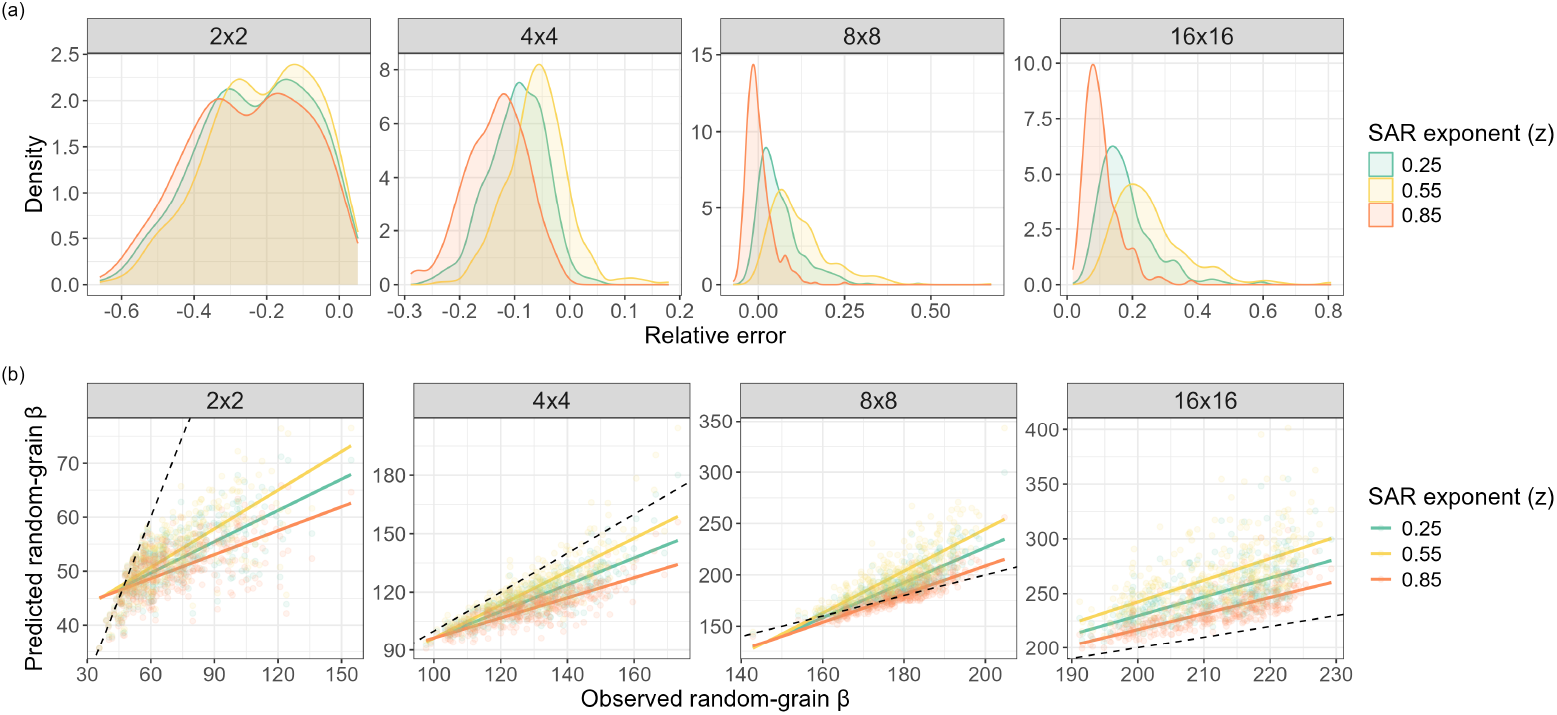
Sensitivity analysis of the Jensen gap approximation for predicting additive beta diversity across randomly sized grains. (a) Distribution of relative error between predicted and observed beta diversity and (b) relationships between predicted and observed beta diversity across different numbers of grains. Predicted beta diversity was calculated using the delta method with three species–area relationships (SAR) exponents (z = 0.25, 0.55, 0.85). Colors represent different values of the SAR exponent (z).

## Footnotes

1 We use *n* for the number of subpopulations here, consistent with the notation of Eq. (1) in the main text and with the partition size *n* introduced in Appendix B.

2 *h* (*x*; *m*_A_) extends continuously to *x* = *m*_A_ (it equals ϕ^′′^(*g* (*x*))2 for some *g* (*x*) between *x* and *m*_A_; see the proof of Theorem 1 in Liao and Berg (2019)). Consequently, provided {*A*_1_, … , *A*_*n*_} ⊂ (*a, b*), the infima and suprema below coincide with the corresponding limits as *x* →*a* or *x*→*b* used in Liao and Berg (2019, Lemma 1).

